# The *period* gene alters daily and seasonal timing in *Ostrinia nubilalis*

**DOI:** 10.1101/2024.11.02.621642

**Authors:** Jacob N. Dayton, Erik B. Dopman

## Abstract

The timing of insects’ daily (feeding, movement) and seasonal (diapause, migration) rhythms affects their population dynamics and distribution. Yet, despite their implications for insect conservation and pest management, the genetic mechanisms underlying variation in timing are poorly understood. Prior research in the European corn borer moth (*Ostrinia nubilalis*) associated ecotype differences in seasonal diapause and daily activity with genetic variation at the circadian clock gene *period* (*per*). Here, we demonstrate that populations with divergent allele frequencies at *per* exhibit differences in daily behavior, seasonal development, and the expression of circadian clock genes. Specifically, later daily activity and shortened diapause were associated with a reduction and delay in the abundance of cycling *per* mRNA. CRISPR/Cas9-mediated mutagenesis revealed that *per* and/or an intact circadian clock network were essential for the appropriate timing of daily behavior and seasonal responsiveness. Furthermore, a reduction of *per* gene dosage in *per* heterozygous mutants (*per* ^-/+^) pleiotropically decreased the diapause incidence, shortened post-diapause development, and delayed the timing of daily behavior, in a manner phenotypically reminiscent of wild-type individuals. Altogether, this combination of observational and experimental research strongly suggests that *per* is a master regulator of biological rhythms and may contribute to the observed life cycle differences between bivoltine (two generation) and univoltine (one generation) *O. nubilalis*.

**Highlights:**

- Natural ecotypes with divergent *period* (*per*) genotypes differ in their daily and seasonal responses to photoperiod
- Later daily activity, reduced diapause incidence, and shorter post-diapause development is associated with reduced *per* mRNA abundance
- *per* is essential for short-day recognition and daily timing
- Reduced *per* gene dosage shortened post-diapause development and delayed locomotor activity

## Introduction

Most plants and animals face strong selective pressures to synchronize their behavioral, physiological, and reproductive cycles with daily and seasonal changes in the environment. Suboptimal timing of these responses can negatively impact individual fitness and population persistence ^1–3^. To counter this, many species have evolved endogenous timing mechanisms that track photoperiod, allowing them to anticipate and respond to rhythmic environmental change.

Insects exhibit photoperiodic diapause, a plastic form of seasonal development comparable to mammalian hibernation. Classical night interruption and resonance experiments have long suggested that the endogenous circadian clock is involved in seasonal photoperiodic time measurement and diapause regulation ^4–7^. The circadian clock relies on transcriptional/translational feedback loops ^8^, and in Lepidoptera, is initiated by the CLOCK (CLK):BMAL1 heterodimer, which drives rhythmic transcription of *period* (*per*), *timeless* (*tim*), and *cryptochrome 2* (*cry2*). In darkness, the resulting TIM, PER, and CRY2 proteins form a complex that translocates into the nucleus to repress CLK:BMAL1-mediated transcription ^9–12^.

Geographic variation in diapause timing has frequently been associated with both allelic variation in clock genes ^13–18^ and differences in circadian timing ^19–21^. Despite these associations, the functional contribution of variation in the circadian clock to the regulation and evolution of diapause remains understudied ^22^. Laboratory manipulations have demonstrated that the circadian clock network is essential for photoperiodic responsiveness in various non-model insects ^23–33^. However, these studies often involved only single populations or genotypes ^22^, which can limit understanding how molecular variation drives population-level divergence. A powerful approach is to combine experimental manipulations with ecologically diverse genotypes ^34–37^, but this approach has not yet widely been applied to insect diapause studies (but see ^13,28,38^).

The European corn borer moth (*Ostrinia nubilalis*) is an agricultural pest that offers an exceptional opportunity to study how genetic variation in circadian clock genes influences behavior and development. Two ecotypes of *O. nubilalis* differ in the number of generations (voltinism) per growing season, largely driven by differences in post-diapause development (PDD) time of overwintering prepupae ^39,40^. Kozak et al. (2019) associated non-coding variants that disrupt an E-box motif within the 5’UTR of the *per* gene with the shorter post-diapause development (PDD) time of individuals from a two-generation (bivoltine) ecotype, which also exhibit longer endogenous circadian periods than their one-generation (univoltine) counterparts. Minor gene expression differences between ecotypes were detected for some clock genes ^41^, but quantitative measurements remain to be estimated for *per* and the rest of the core clock gene network. These findings suggest that variation in *per* genotype and function might contribute to the phenotypic variation in both daily and seasonal traits ^42^.

Here, we integrate behavioral assays, transcriptome profiling, and targeted genetic/pharmacological manipulations to investigate *per*’s function in both biological rhythms. Our results demonstrate that the circadian clock gene *per* is essential for timekeeping and reductions to *per* gene dosage directly alter both daily and seasonal rhythms. Based on these laboratory results, we hypothesize that *per* genotype-specific differences in *per* transcript abundance may contribute to the ecotype divergence in *O. nubilalis*, offering new insights into how genetic variation in clock genes may shape adaptive responses in insects.

## Results and Discussion

### *Ostrinia nubilalis* ecotypes differ in timing

The relationship between seasonal and daily timing was explored by measuring the prepupal diapause response and overt activity rhythms of univoltine and bivoltine *O. nubilalis* ecotypes with divergent allele frequencies at *period* (*per*). *O. nubilalis* were responsive to subtle changes in photoperiod and reproduced diapause phenotypes previously documented in the field ^39,43^ and laboratory ^42,44^. As daylength decreased, diapause incidence increased, and fewer individuals continuously developed into pupae (Fig. 1A). Genetic variation in the response to photoperiod was reflected by the statistically significant effects of ecotype (i.e., genetic background) and the interaction between ecotype and photoperiod on both diapause incidence and post-diapause development (PDD) time (Fig. 1A-1B; see Fig. S1). The bivoltine ecotype exhibited lower diapause incidence in longer photoperiods (Fig. 1A) and a shorter PDD time after transfer to diapause-breaking conditions (Fig. 1B; see Fig. S1).

**Fig. 1.**
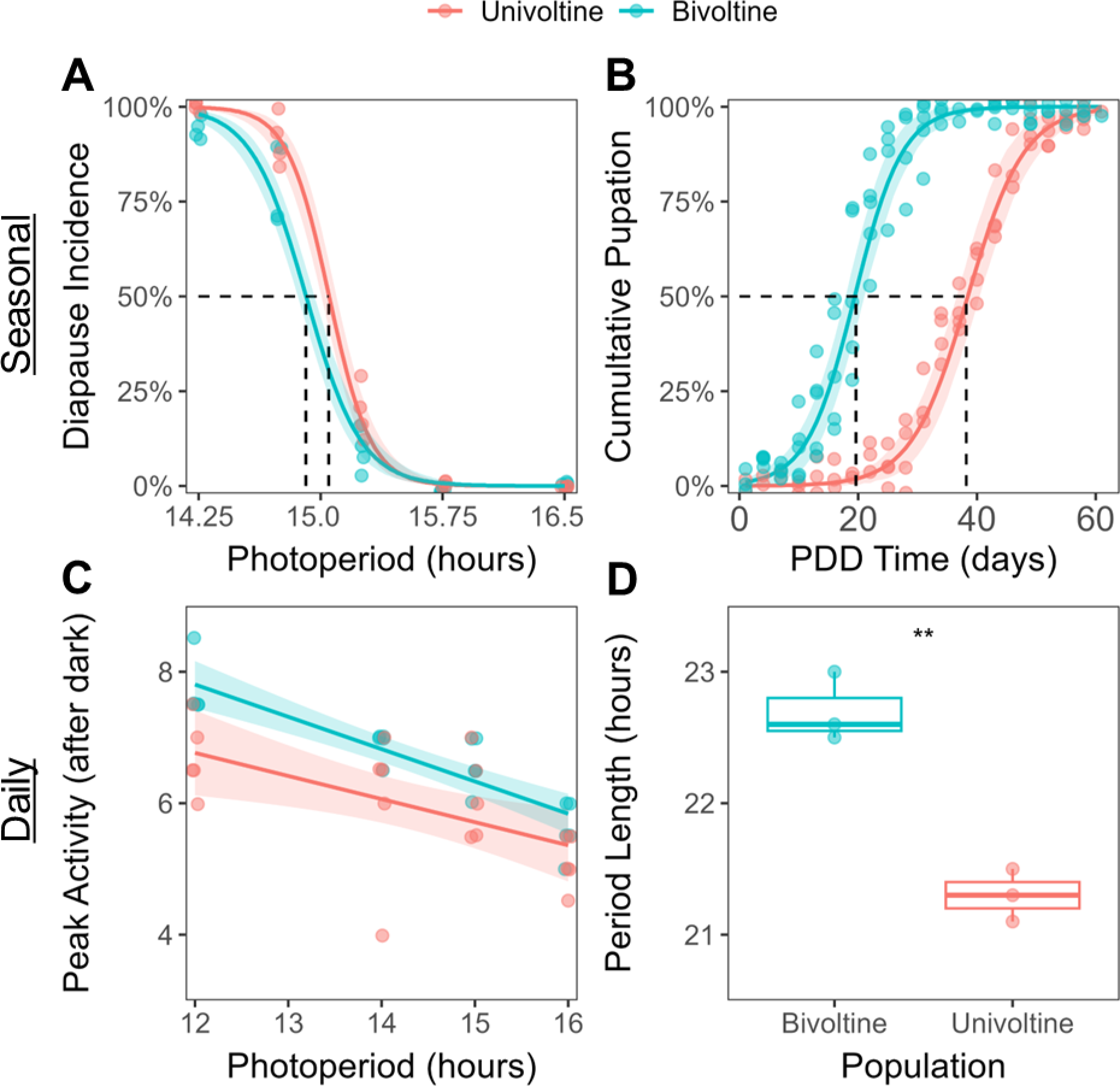
*O. nubilalis* ecotypes differ in seasonal and daily rhythms. (A) Diapause induction in larvae exposed to various light-dark (LD) photoperiods at 23.5°C. Diapause was determined as the failure to pupate by day 35. Photoperiod (LR χ^2^ = 1108, *df* = 1, *P* = <2.2*10^-16^), ecotype (LR χ^2^ = 15.55, *df* = 1, *P* = 8.05e-5), and their interaction (LR χ^2^ = 4.99, *df* = 1, *P* = 0.026) significantly predicted diapause incidence (binomial GLM, logit link). Dashed lines denote the Critical Daylength (CDL), where the probability for diapause is 50%, for univoltine (15.05 L) and bivoltine (14.89 L) populations. (B) Diapausing larvae from 14.25L:9.75D were transferred to 16L:8D to track post-diapause development (PDD) time to pupation. PDD time was significantly affected by photoperiod (binomial GLM; χ**^2^** = 123, *df* = 1, *P* < 2.2*10^-16^), ecotype (χ**^2^** = 1810, *df* = 2, *P* < 2.2*10^-16^), and their interaction (χ**^2^**= 125, *df* = 2, *P* < 2.2*10^-16^). Bivoltine individuals completed PDD ∼20 days earlier than 50% of univoltine individuals (38.5 days). See Fig. S1 for results from other diapause-inducing photoperiods. (C) Across LD photoperiods, the phase of peak locomotor activity was significantly affected by daylength (F = 41.5, *df* = 1, *P* = 1.78*10^-7^) and ecotype (F = 14.0, df = 1, *P* < 0.001), but not their interaction (*P* = 0.291). (D) In continuous darkness (DD), the free-running period length of bivoltine males was 1.4 hr longer (95% CI: 0.85−1.95 hr) than univoltine males (*t* = 7.31, *df* = 3.72, *P* = 0.002). Shading denotes 95% confidence intervals. Asterisks denote P < 0.01 (**).

Highlighting possible connections between the circadian clock and seasonal timing, a shifted phase (Φ) in the daily timing of peak locomotor activity (Fig. 1C) accompanied these ecotype differences in seasonal diapause. Bivoltine males were active 0.72 hr (Φ, 95% CI: 0.33−1.12 hr) later than univoltine males (normal distribution; *t* = 3.74, *df* = 37, *P* = 0.006; Fig. 1C). Specifically, in *D. melanogaster*, an advanced phase of activity (LD) and a shorter free-running period (DD) reflect earlier PER-mediated transcriptional repression, driven by increased *per* gene dosage ^45–47^, elevated transcription of *per* ^46,48,49^ and/or altered phospho-regulation of PER protein stability/abundance and nuclear localization ^50–54^. Accordingly, the later activity phase of bivoltine males was consistent with their longer free-running period in DD (Fig. 1D). Although some components of the Lepidopteran circadian clock differ from *D. melanogaster ^8^*, we hypothesized that these behavioral phenotypes in bivoltine *O. nubilalis* may similarly represent a delay in PER repression, driven by reduced *per* gene expression and/or delayed PER protein accumulation during the evening.

### *O. nubilalis* ecotypes differ in clock gene expression

To explore potential transcriptional mechanisms underlying ecotype differences, we leveraged 3’-end RNA sequencing ^55,56^ to quantify cycling gene expression in *O. nubilalis* brains, the site of the central circadian pacemaker and photoperiodic timer ^7^. Across a 12L:12D cycle, RNA was isolated from replicate pools of fifth instar diapause-destined brains at 3-hr intervals, starting from the scotophase of day 4 through the photophase of day 5. Although cosinor regression identified 1,230 rhythmic genes (12% of expressed genes), with 773 that were differentially rhythmic ^57^ between ecotypes (*q_DR_* < 0.05), we focused our investigation on the expression of the core circadian clock genes in Lepidoptera ^8^(Fig. 5B).

Within the *O. nubilalis* brain, ecotypes differed in their gene expression profiles for core components of the circadian clock gene network. Later evening activity of bivoltine individuals (Figs. 1C, 1D) was mirrored at the transcriptional level by a 1.4 hr later mean phase (Φ) of cyclic expression of *per, cryptochrome 2* (*cry2*), and *vrille* (*vri*) genes (95% CI: 1.0−1.7 hr; *t* = 17.0, *df* = 2, *P* < 0.003; Fig. 2), all of which were significantly rhythmic in both ecotypes (Fig. 2). Pairwise comparison of these genes revealed significantly lower rhythm-adjusted mean (mesor) expression of *per* (*P* = 0.027) and higher mesor expression of *vri* (*P* = 0.001) in bivoltine individuals (Fig. 2). For the six genes lacking cross-ecotype rhythmicity, *clk* was downregulated (log_2_FC = -0.56, q = 0.004) and both *cyc* (log_2_FC = -1.06, q = 0.003) and *tim* (log_2_FC = -0.46, q = 0.002) were upregulated in the bivoltine ecotype.

**Fig. 2.**
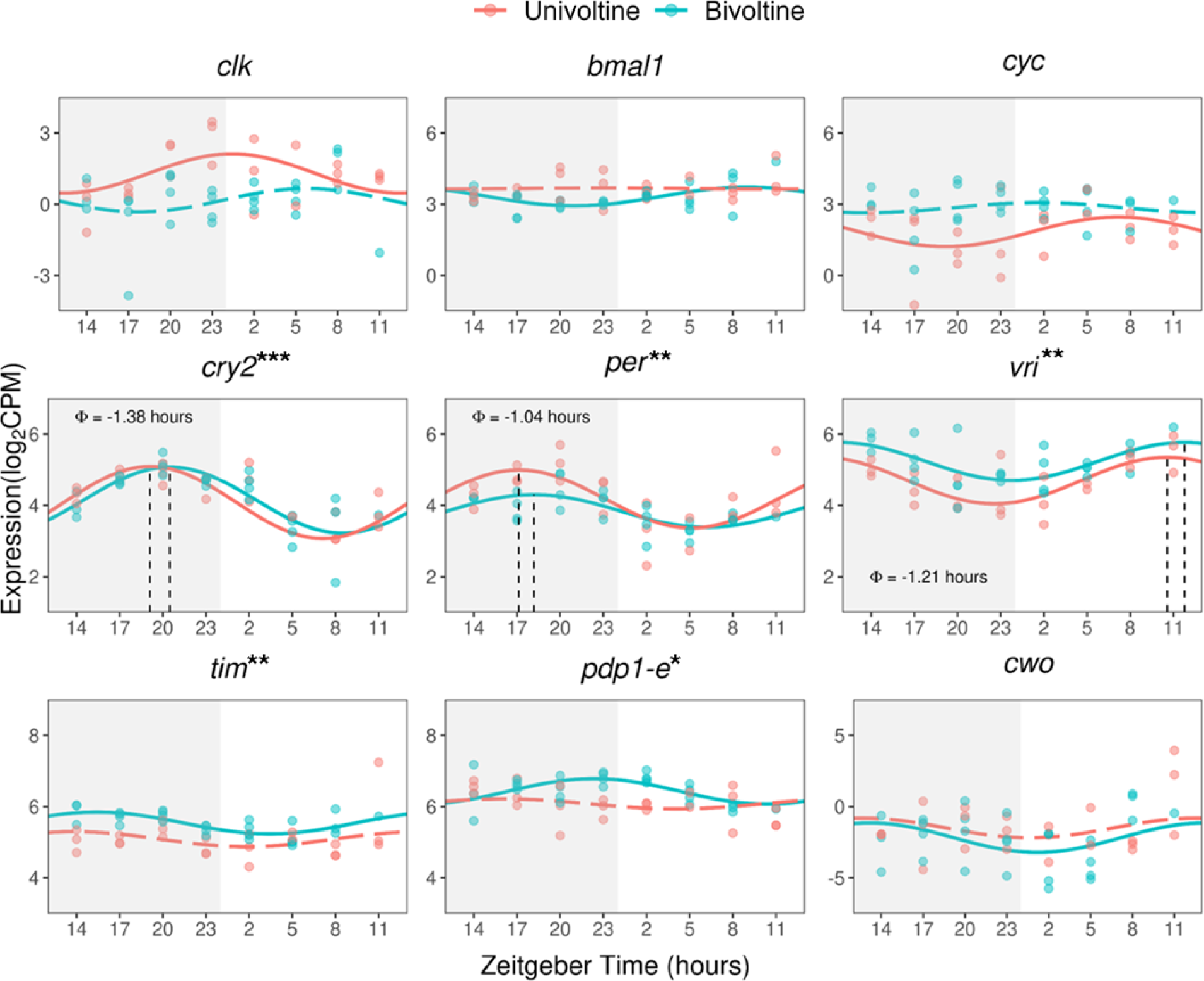
*O. nubilalis* ecotypes differ in rhythmic expression of core circadian clock genes in the brain. Larvae (12L:12D, 23.5°C) were synchronized at molt into the final 5th instar (day 0). Pools of 3-4 brains were dissected every three hours from day 4 (ZT14) through day 5 (ZT11). Darkness is shaded (ZT12-24). Across ecotypes, evidence for significant rhythmicity (cycling) in gene expression was evaluated by cosinor regression and adjusted for multiple comparisons (B-H method, *q* value). Asterisks denote *q*_rhy_ < 0.10 (*), *q*_rhy_ < 0.05 (**), and *q*_rhy_ < 0.01 (***). Genes that are rhythmic within ecotypes (*P*_rhy_ < 0.05) are illustrated with solid fitted cosine-regression lines; dotted lines indicate no significant cosine fit curve (*P*_rhy_ ≥ 0.05). Comparisons of rhythmic genes between ecotypes revealed significant differences in the mesor (*per*, *P* = 0.027; *vri*, *P* = 0.001) and consistent differences in phase (Φ). *clk*, *clock*; *bmal1*, *brain and muscle Arntl-like 1*; *cyc*, *cycle*; *cry2*, *cryptochrome 2*; *per*, *period*; *vri*, *vrille*; *tim*, *timeless*; *pdp1-e*, *par domain protein 1*; *cwo*, *clockwork orange*. *Drosophila*-like *cryptochrome 1* is not displayed and was not rhythmic (*q*_rhy_ > 0.10).

Based on the function of circadian clock genes in *D. melanogaster* ^58^ and Lepidoptera *^8^*, the reduced expression of *clk* and *per* ^9,10,12^, paired with upregulated *vri ^59,60^*, jointly suggested that the bivoltine population’s later activity (Fig. 1C) and longer free-running period (Fig. 1D) could be a consequence of reduced evening accumulation of PER and a dose-dependent delay in PER-mediated repression (Fig. 5A-5B). Although the connection between differences in the circadian clock network and seasonal timing is less clear in the literature, the prior association between ecotype variation in diapause responses and *per* genotype^42^ suggested that amplifying effects of reduced *per* (Fig. 2) across days may also contribute to the lower diapause propensity (Fig. 1A) and shorter PDD time of bivoltine individuals (Fig. 1B).

### Reduced *per* gene dosage confers bivoltine-like responses

Given that the shorter PDD time of bivoltine *O. nubilalis* (Fig. 1) is associated with genetic variation in the *per* 5’ UTR ^42^ and reduced *per* gene expression (Fig. 2), we leveraged CRISPR/Cas9 to investigate whether *per/*PER abundance directly alters daily behavior and seasonal diapause. Since the number of wild-type *per* gene copies in *D. melanogaster* is inversely correlated with the free-running period length ^45–47,61^ and *per* mRNA abundance ^46^, we predicted that reducing *per* gene dosage would similarly decrease the abundance of *per*/PER in *O. nubilalis* and resemble bivoltine-like phenotypes. We generated germline mutants bearing frameshift mutations in *O. nubilalis per* exon four ^62^, upstream of the predicted Per-Arnt-Sim (PAS) domain. Consistent with reports from other insects ^30,31^, *per* hemizygous female mutants (*per* ^-^/W) were unresponsive to a diapause-inducing photoperiod (see Fig. S2A), and they exhibited arrhythmic adult eclosion behavior (see Fig. S2B), indicative of a dysfunctional circadian clock ^11,30,31,63^. In the absence of an *O. nubilalis*-specific PER antibody, these defects suggested that frameshifts at this sgRNA target site resulted in a true loss-of-function allele.

In contrast, *per* heterozygous mutants (*per ^-/+^*) were still responsive to photoperiod and exhibited nearly 100% diapause incidence in 12L:12D (Fig. 3A). However, when larvae in a diapause-inductive photoperiod (12L:12D) were exposed to a late-night light pulse, known to accelerate PER degradation ^10,64^, diapause incidence was drastically reduced (Fig. 3A). Resembling the lower diapause incidence (Fig. 1A) and elevated photosensitivity of bivoltine wild-type *O. nubilalis* (see Fig. S3), *per* heterozygous mutants (*per ^-/+^*) with only one functional *per* gene copy were more sensitive to diapause-averting night interruptions (Fig. 3A). This similarity between laboratory mutants and wild-type bivoltine individuals suggested that *per* (and PER repression) may directly be involved in measuring night length, the causal factor determining diapause incidence in *O. nubilalis ^4,65^* and many other insects ^6,7^.

**Fig. 3.**
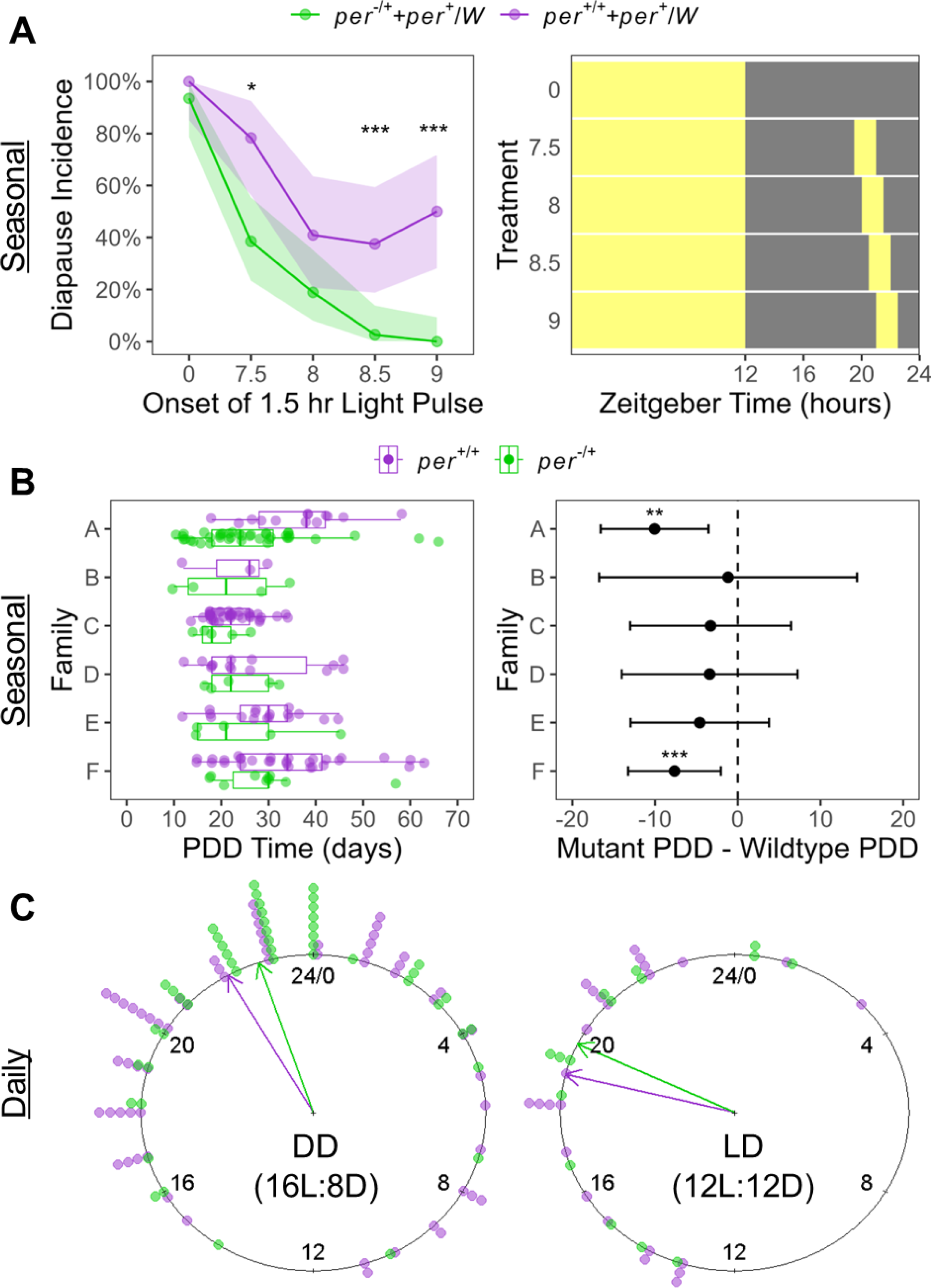
Reduced *per* gene dosage pleiotropically modifies seasonal development and daily behavior. (A) Diapause induction of larvae reared in 12L:12D and exposed to a 1.5 hr night-interruption light pulse (no pulse = 0). Diapause incidence was significantly affected by photoperiod treatment (χ^2^ = 131.3, *df* = 4, *P* < 2.2*10^-16^), genetic background (χ^2^ = 44.5, *df* = 1, *P* < 2.57*10^-11^), and their interaction (χ^2^ = 11.2, *df* = 4, *P* = 0.025). Significant differences were evaluated by a two-sample proportion test. Shading denotes 95% confidence intervals.(B, left) Diapausing larvae (12L:12D) were transferred to 16L:8D to track post-diapause development (PDD) time. Each family contained paired *per* homozygous wild-type (*per ^+/+^*) and *per* heterozygous mutant (*per ^-/+^*) siblings, genotyped by allele-specific PCR or amplicon sequencing. (B, right) Intra-family differences in mean PDD time between heterozygous mutants and wild-type siblings. Only *per* gene dosage (two-way ANOVA: F = 15.8, *df* = 1, *P* < 0.001) and family (F = 5.51, *df* = 5, *P* < 0.001), but not their interaction (F = 0.54, *df* = 5, *P* = 0.742), significantly predicted PDD time. Averaged across families, *per* heterozygous mutants exhibited a 6.6 day shorter PDD time (95% CI: 3.3−9.9 days, *t* = 4.0, *P* < 0.001). (C) Eclosion of adults from their pupal case in DD (Left) and LD (Right) was binned into 1 hr intervals and pooled across families. (C, left) Pupae were kept in 16L:8D and transferred to continuous darkness (DD) seven days after pupating. (C, right) Pupae were kept in 12L:12D. The lights turned on at Zeitgeber Time 0/24. Arrows denote the circular mean eclosion time, which was delayed in *per* heterozygous mutants in both DD (Φ = 0.76 hr; Watson’s Two-sample U^2^= 0.215, *P* < 0.05) and LD (Φ = 0.77 hr; U^2^ = 0.057, *P* > 0.10). Asterisks denote *P* < 0.05 (*), *P* < 0.01 (**), *P* < 0.001 (***).

To conservatively isolate the effect of *per* gene dosage from any differences between families, arising from genetic background or a family-specific microenvironment, we reared *per* heterozygous mutants (*per ^-/+^*) alongside their respective wild-type (*per ^+/+^*) siblings in 12L:12D. Diapausing larvae/prepupae were transferred to 16L:8D to terminate diapause and track PDD to pupation. Indicating a direct link between *per* gene dosage and seasonal timing, PDD time was significantly predicted by *per* gene dosage and family (Fig. 3B), and *per* heterozygous mutants exhibited a 7-day shorter PDD time on average (Fig. 3B).

A complementary experiment investigated the effect of different diapause-inducing photoperiods (15L:9D vs. 12L:12D hr) and *per* gene dosage on PDD time. The absence of an interaction between *per* gene dosage and photoperiod (F = 0.38, *df* = 1, *P* = 0.539) suggested that photoperiod and *per* independently exert their influence on PDD time (see Fig. S4).

To investigate whether a reduction in *per* gene dosage simultaneously altered the circadian clock network, the daily eclosion of adults was separately monitored in LD or after transfer to DD. Consistent with the period lengthening reported in *Drosophila* with reduced *per* gene dosage^45^, the circular mean eclosion time of *per* heterozygous mutants (*per ^-/+^*) was significantly delayed in DD (Φ = 0.76 hr) and similarly affected in LD (Φ = 0.77 hr; Fig. 3C). Altogether, the analysis of *per* heterozygous mutants revealed that *per* gene dosage (and unmeasured *per/*PER abundance) directly altered both daily and seasonal timing.

### Pharmacological period lengthener reduces diapause incidence

Mutations in a single clock gene, like *per*, cannot easily be isolated from any effect on the entire circadian clock module ^25,28,66^. Therefore, to gather additional evidence that delayed PER repression in the circadian clock alters seasonal timing, we performed pharmacological manipulations of wild-type larvae. In *D. melanogaster* and mammals, lithium lengthens the free-running period of the circadian clock ^67–69^ by inhibiting GSK-3β/SGG-mediated phosphorylation, which can delay nuclear localization of PER ^53^. Based on the longer free-running period and reduced diapause incidence of bivoltine *O. nubilalis* (Fig. 1), we hypothesized that dietary lithium supplementation would decrease diapause incidence in a dose-dependent manner reminiscent of reduced *per* gene dosage in *per* heterozygous mutants (Fig. 3A). Consistent with this prediction, increasing concentrations of lithium decreased diapause incidence in both ecotypes (β = -0.027, z = -2.50, *P* = 0.012; Fig. 4). This effect was more pronounced in the bivoltine population (β = -0.029, z = 1.87, *P* = 0.062; Fig. 4), and may reflect the greater slope change near a population’s critical daylength (Fig. 1A).

**Fig. 4.**
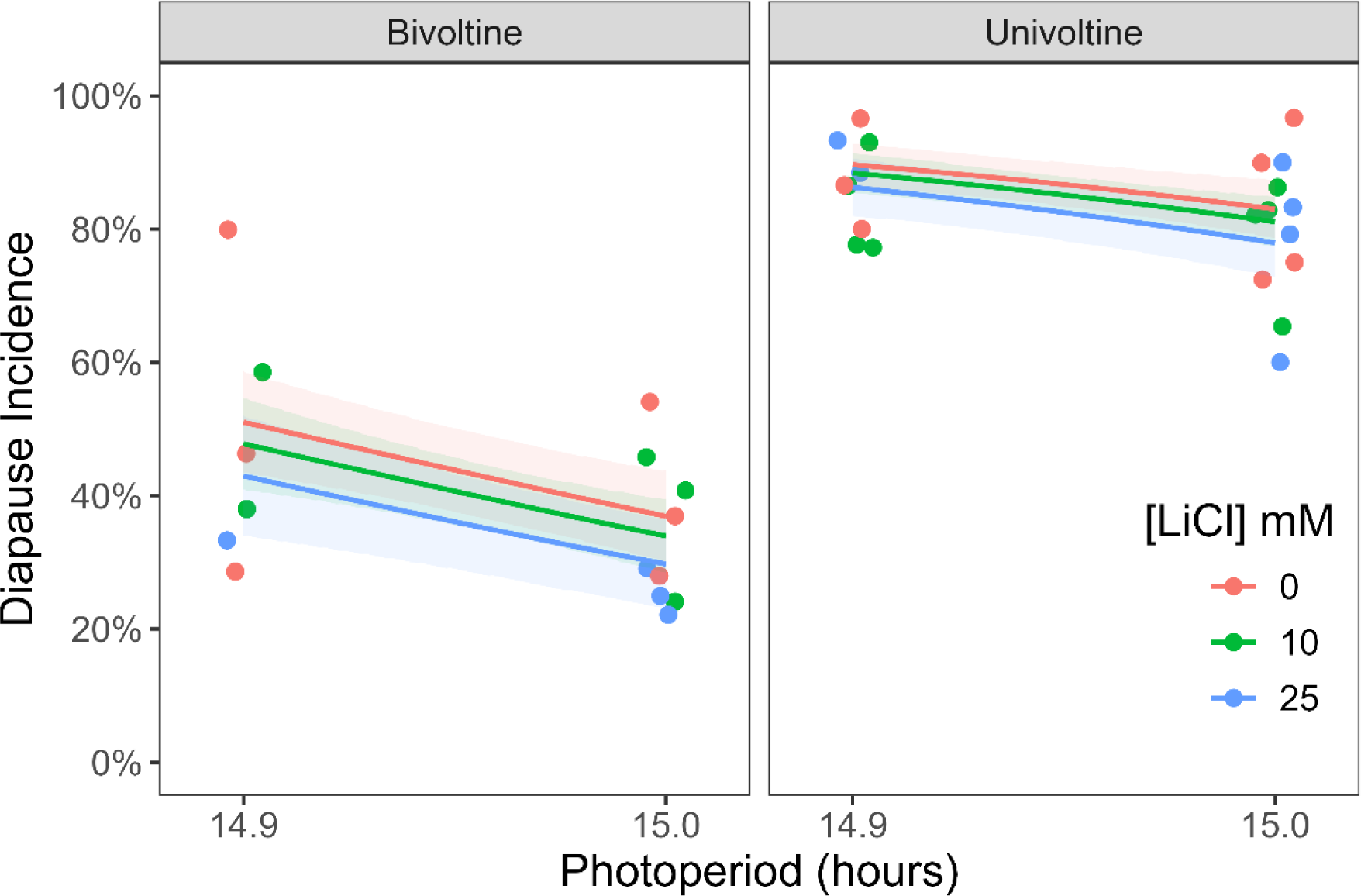
Lithium reduces diapause incidence in *O. nubilalis*. Groups of larvae from bivoltine and univoltine ecotypes were reared in diapause inducing conditions (14.9L:9.1D, 15L:9D). From the onset of the fifth instar, larvae were isolated and fed a diet containing 0, 10, or 25mM of LiCl. The effects of lithium concentration, photoperiod, and background on diapause incidence were analyzed by a binomial GLM using a logit link. Lines denote model predicted diapause incidence for each combination of fixed effects. Diapause incidence was significantly affected by ecotype (binomial GLM; χ^2^ = 216, *df* = 1, *P* = 2.2*10-16) and photoperiod (χ^2^ = 12.45, *df* = 1, *P* < 0.001), with evidence for a lesser effect by lithium (χ^2^ = 2.914, *df* = 1, *P* = 0.088) and the interaction between ecotype and concentration (χ^2^ = 3.53, *df* = 1, *P* = 0.060). Each point represents a replicate group (28-30 larvae). Shading denotes 95% confidence intervals.

### A possible mechanism for biological rhythm

Overall, we demonstrate that multivariate trait divergence in daily and seasonal timing (Fig. 1) of *O. nubilalis* ecotypes is associated with *per* genotype ^42^ and altered expression of cycling *per* mRNA (Fig. 2). Laboratory manipulations predicted to delay PER repression, either by reducing wild-type *per* gene dosage (Fig. 3) or lithium treatment (Fig. 4), induced changes resembling wild-type bivoltine *O. nubilalis*. Although we have not functionally validated the influence of naturally-occurring *per* alleles, this combination of observational and experimental research implicates the hypothesis that variation in the circadian clock network directly modifies both daily and seasonal traits. In the case of *O. nubilalis*, *period* is predicted to be a master regulator of daily and seasonal rhythms: reduced *per* mRNA (and PER) abundance during the evening delays daily activity and contributes to a less intense diapause response (Fig. 5). Moreover, finding that a conserved period lengthener (lithium) also reduces diapause incidence suggests that diverse manipulations of circadian clock properties (e.g., amplitude, phase) can also predictably modify seasonal timing. By extension, manipulations ^49,70^ or environmental conditions ^28,71–73^ that increase *per* mRNA and advance evening accumulation of PER may confer earlier activity and a more intense diapause response (Fig. 5).

**Fig. 5.**
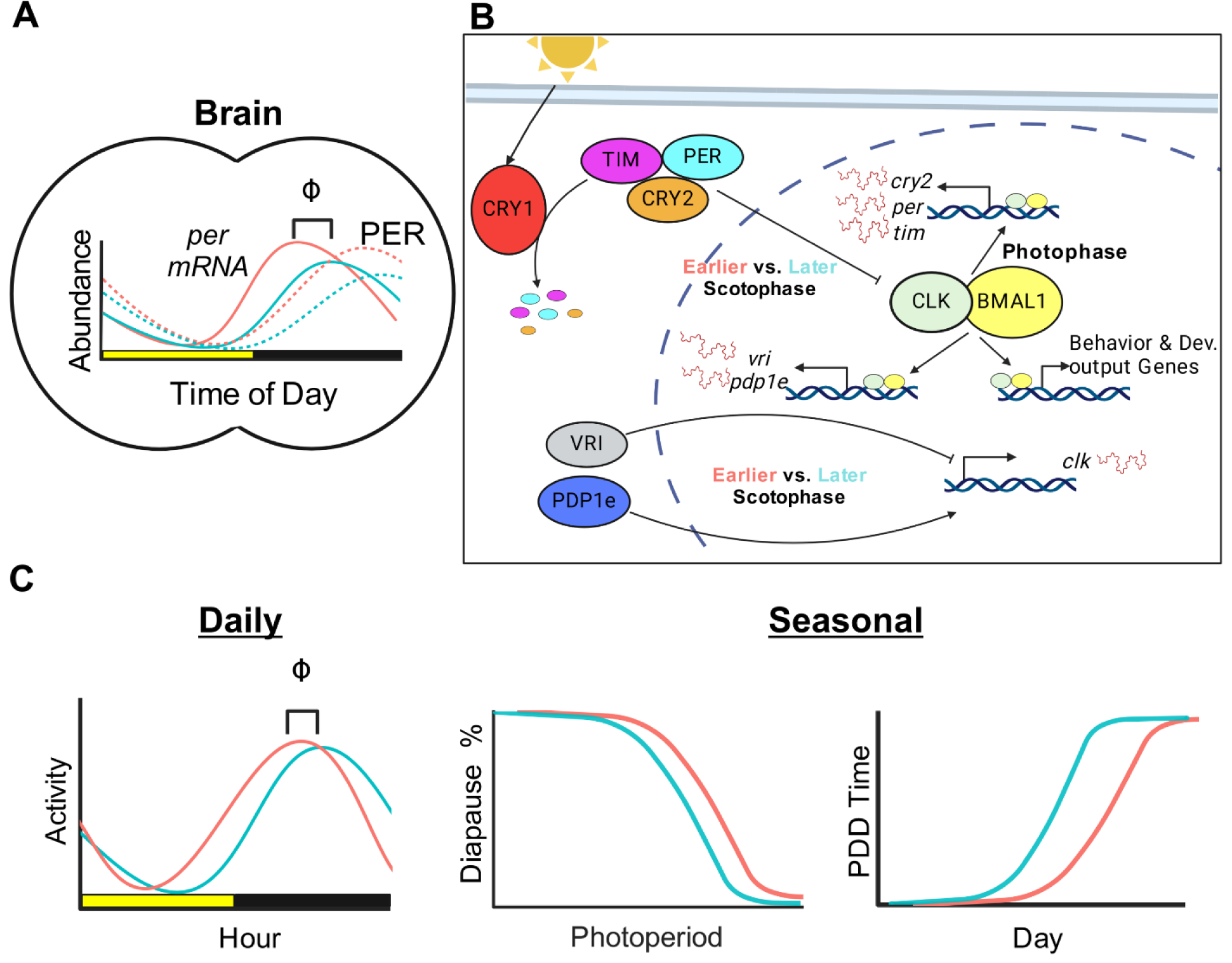
*per* transcript abundance alters daily and seasonal rhythms in *O. nubilalis*. (A) Variation in the phase (Φ) or amplitude of *per* mRNA abundance, and predicted PER protein accumulation, across a 24-hr day is associated with differences in daily behavior and seasonal development of univoltine (red) and bivoltine (blue) individuals. (B) In a simplified model of the Lepidopteran circadian clock network^8^, we hypothesize that earlier vs. later repression of CLK:BMAL1 in the evening (scotophase) pleiotropically alters both daily and seasonal rhythms of wild-type individuals. (C) Experimental reductions in *per* gene dosage (blue) delay the phase of daily behavior, decrease photoperiodic diapause incidence, and shorten post-diapause development (PDD).

Nevertheless, we note limitations to our study that will require further work. Although the relationship between *per* mRNA levels and the abundance of its cognate protein are tightly associated in *D. melanogaster ^54^*, we have not yet quantified in *O. nubilalis* how *per* gene expression directly alters PER protein abundance. Secondly, although empirical associations between nuclear localization of the repressive complex, requisite for PER repression ^9^, and evening photosensitivity have been documented in *D. melanogaster ^74^*, PER immunostaining is needed to resolve the neuroanatomical structure of the clock and to identify the onset of repression in *O. nubilalis*. Similarly, coimmunoprecipitation (CoIP) of PER with CLK:BMAL1 or chromatin (ChIP) could respectively inform when PER sequesters CLK from DNA ^12,75^ and reveal transcriptional targets mediating different daily and seasonal responses. These targets and their downstream signaling pathways ^33^ could also be nominated from further analysis of the transcriptomic resources developed here.

Although *per* was the focus of this manuscript, Kozak et al. (2019) also associated differences in seasonal timing with variation in the *Pigment dispersing factor receptor* (*Pdfr*) gene. *Pdfr* was hypostatic to allelic changes at *per* and explained less phenotypic variance in PDD time. However, considering that *Pdfr* encodes the receptor for the neuropeptide PDF ^76^, and PDF/PDFR signaling interacts with PER ^64,77^ and TIM ^78^ to adjust the phase and amplitude of clock neurons under long-day photoperiods ^77,79^, further exploration of its contribution to daily and seasonal timing in *O. nubilalis* is warranted. Already studies in *Culex pipiens* mosquitoes and *D. melanogaster* have shown that PDF signaling is essential for long day responses, and loss of PDF caused insects reared in long day photoperiods to enter a diapause-like state ^25,80–82^. In contrast, PDF knockout prevented diapause induction in *Pyrrhocoris apterus* and *Plautia stali* individuals exposed to short days ^83–85^. Therefore, an important question concerns how allelic variation at *Pdfr* may alter PER oscillations and contribute to photoperiodic responses in *O. nubilalis*.

Finally, it will be important to investigate how transferable these results are across populations and species. For example, the hormonal control of diapause regulation is relatively consistent across species that diapause in the same life-cycle stage ^7^. However, relationships between diapause timing and either variation in circadian clock properties ^20^ or in the response to clock gene manipulation ^33^ are not the same across species. Despite this, continued studies of diapausing insects, spanning diapause strategies, life-cycle stages, and mechanisms of photoperiodic time measurement (*i.e.,* night-length, daylength), might reveal commonalities in how changes to the circadian clock’s function drive seasonal adaptation.

In summary, we leverage empirical studies of diverse populations with laboratory manipulations to provide evidence that differences in *per* mRNA abundance contribute to the key differences in seasonal timing of univoltine and bivoltine *O. nubilalis* ecotypes. While we predict that reduced *per* mRNA levels delays evening PER accumulation and PER repression of the circadian clock, the specific mechanisms and their downstream targets require further study. Regardless, these results implicate a link between variation in the circadian clock and adaptive differences in seasonal timing.

## Data and Code Availability

Raw sequencing data are being submitted to GEO and a GEO:accession number will be provided when assigned. Processed data and original code are publicly available on GitHub: https://github.com/daytonjn/period_ms_2024 and will be accessible at Zenodo via a DOI before publication. Any additional information required to reanalyze the data reported in this paper is available upon request from the lead contact, Jacob Dayton.

## Acknowledgements

We thank K. McLaughlin for access to critical microinjection equipment, A. Balikian, R. Klusza, A. Murray, J. Paul, and T. Tran and for assistance in the lab, E. Saint-Denis for assisting with tissue dissections, and S. Mirkin, B. Trimmer, and M. Meuti for thoughtful feedback on an early version of the manuscript. The authors acknowledge the Tufts University High-Performance Computing Cluster https://it.tufts.edu/high-performance-computing), which was utilized for the research reported in this manuscript. Figure 5 was generated on BioRender with a license to J.N.D.

## Funding Information

E.B.D. acknowledges support from the National Science Foundation (Award #2416175) and Tufts University.

## Author Contributions

Conceptualization, J.N.D., E.B.D.; Methodology, Investigation, Formal analysis, Data curation, Writing−initial draft, J.N.D.; Writing−review, J.N.D., E.B.D.; Funding acquisition, E.B.D.

## Declaration of Interests

The authors declare no competing interests.

## Supplemental Information

Document S1. Figures S1-S4 and Tables S1-S2.

## STAR Methods

### Insect stocks

Univoltine and bivoltine European corn borers (ECB; *Ostrinia nubilalis*) were collected from laboratory populations maintained at Tufts University (Medford, MA). These colonies were originally derived from insects collected in corn stubble in New York, USA and New Hampshire, USA. These colonies have repeatedly been selected for univoltine vs. bivoltine PDD-time^42^ and exhibit divergent allele frequencies at *per*. As described previously^62^, *O. nubilalis* larvae were reared on artificial corn borer diet (Southland Products, USA) and maintained in an environmental room at 16L:8D (25.5°C; 55% RH).

### Phenotyping diapause in wild-type individuals

To induce diapause, eggs were suspended over diet and reared in climate-controlled incubators for 12L:12D hr at 23.5°C (50% RH). Fourteen days after hatching, larvae were transferred into individual diet-containing 1.25 oz plastic souffle cups and transferred to various photoperiods differing in the duration of photophase. Photoperiodic response curves for diapause incidence were derived from individuals exposed to 14.25L:9.75D, 14.75L:9.25D, 15.25L:8.75D, and 16.5L:7.5D hr. Larvae who failed to pupate before Day 35 were classified as diapausing ^86,87^; by this day, the cumulative pupation curve has already leveled off. On Day 35, larvae were transferred to 16L:8D to artificially terminate diapause ^42^. To score PDD-time, individuals were checked every two days for pupation.

### Phenotyping diapause in response to night-interruption by light

Experiments predominantly utilized bivoltine and univoltine larvae. To test the influence of *per* gene dosage on diapause incidence, a bivoltine *per* hemizygous mutant female (*per ^-^/W*) was crossed with a wild-type univoltine male to generate *per* heterozygous mutants (*per ^-/+^*) and wild-type females (*per ^+^/W*). Larvae were initially reared from hatching in a photoperiod known to induce diapause (12L:12D hr at 26°C). After 14 days, larvae were isolated into individual cups with diet and transferred into various night interruption treatments to determine the position of the photoinducible phase ^6,88^. Night interruption treatments were conducted in opaque boxes with overhead warm white LED strip lights. LED lights were programmed with either BN-LINK or myTouchSmart digital outlet timers. Every treatment box contained 48 individuals. A black towel covered boxes to prevent any light from escaping.

Night interruption experiments consisted of a 12L:*x*D:*y*:(12-*x-y*)L hr photoperiod (T=24), with *x* denoting the onset of the interruption light and *y* denoting the duration of the light pulse ^89^. The larvae were maintained in night interruption treatments for 16 days. Individuals who failed to pupate by day 32 were considered in diapause. Larvae that did not survive the length of the experiment were not included in the results. One bivoltine family was completely non-diapausing and pupated under all photoperiods; these were removed from analyses. The effects of genetic background, night interruption timing (factor), and their interaction on diapause incidence were modeled by a binomial GLM with a logit link. Significant differences in diapause incidence were evaluated by two-sample proportion tests.

### Phenotyping diapause in *per* mutant families

Offspring of different *per* heterozygous mutant males and wild-type females were exposed to either 12L:12D hr (Families A-F) or 15L:9D hr (Families E-F) at 23.5°C. After 35-42 days in inducing conditions, diapausing larvae were transferred to 18°C for 7 days and then stored at 4°C for 45 days. To terminate diapause and promote post-diapause development, larvae were transferred to 16L:8D and checked every two days for pupation. Quiescent larvae that rapidly pupated within eight days were excluded from analyses. The experiment was initially run in 2022 (Families A-D) and repeated in 2024 (Families E-F).

### Effect of pharmacologic period-lengthening on diapause induction

European corn borer larvae were reared in 14.9L:9.1D and 15L:9D hr (23.5°C). Upon head-capsule slippage preceding molt to 5^th^ instar, larvae were removed from their artificial diet (Southland Products), transferred into one of two photoperiodic conditions previously shown to induce diapause in 50% of the population (14.9:9.1 LD and 15:9 LD hr), and placed onto fresh diet containing an additive. Larvae who failed to pupate by day 35 were considered in diapause. At this temperature, the cumulative pupation curve has already leveled off in a control population. The experiment was separately conducted with bivoltine (Fall 2020) and univoltine (Spring 2021) individuals. Across both experiments, 3-4 replicate groups of 30 larvae were exposed to various LiCl concentrations (0mM, 10mM, 25mM LiCl). The effects of pharmacologic treatment and photoperiod on diapause incidence were determined by a binomial regression with a logit link. Anova() in the *car* package evaluated significance for effects and interactions.

### Phenotyping eclosion

Eclosion was monitored by the Raspberry Pi-based imaging Locomotor Activity Monitor ^90^, as previously described in Dayton et al. (2024). The only modifications were that in 2022, pupae from Families A-D were transferred into continuous darkness seven days after pupation. In 2024, pupae from Families E-F were transferred upon pupation into 12L:12D and monitored for eclosion in LD.

### Phenotyping locomotor activity

The imaging Locomotor Activity Monitor (iLAM)^90^ quantified locomotor activity of adult males exposed to entraining (LD) and free-running (DD) conditions. Activity of three replicate flight cages containing four to five males each was recorded for 4-5 days in an enclosed room at 23°C (40% RH). For LD experiments, paired flight cages were exposed to 12L:12D, 14L:10D, 15L:9D, or 16L:8D hr photoperiods by connecting overhead LED lights (3000K Warm White, Pautix) to programmed outlet timers (BN-LINK). Individual photoperiod conditions were isolated from each other by multiple blackout curtains. The time that lights turned on was synchronized across cages and denoted as Zeitgeber Time 0 (ZT0). For circadian phenotyping in DD, moths were acclimated in flight cages for one day in 16L:8D before exposure to DD.

Image segmentation and movement quantification utilized the *iLAMtools* wrapper functions ^90^ for *imager ^91^*. All analyses occurred in the *Rethomics ^92^* framework. Activity in LD was smoothed with a Butterworth filter and *pracma ^93^* identified the daily timing of peak activity. Free-running period length (DD) was estimated by chi-squared periodogram.

### RNA library preparation and sequencing

Diapause-destined *O. nubilalis* larvae (12L:12D, 23.5°C) from univoltine and bivoltine ecotypes were staged at molt into the final 5^th^ instar (day 0). Beginning on the fourth day of the fifth instar, four replicate pools of 3-4 brains each were isolated every three hours from Day 4 (ZT14) through Day 5 (ZT11). Larvae were decapitated and brains were dissected in DNA/RNA Shield (Zymo Research, R110050) reagent and frozen at -20°C. To avoid batch effects due to differences in RNA purification efficiencies, all homogenized samples were purified in parallel using the Quick-RNA MicroPrep Kit (Zymo Research, R1050). Total RNA was eluted in water, quantified with the NanoDrop and diluted to ∼20ng/uL. To avoid batch effects in cDNA amplification and library preparation, we prepared a high-throughput Mercurius BRB-seq 3’-mRNA library (Alithea Genomics, 10813) following manufacturer’s instructions, with ∼200ng RNA from each sample. Cleanup and concentration steps used HighPrep PCR magnetic beads (MagBio Genomics, AC-60005). Tagmentation with pre-loaded adapters used 50ng of cDNA incubated for 8 minutes at 55°C. Only 12 cycles were used for library indexing and amplification. Sequencing was performed by the Tufts University Core Facility Genomics and used the Illumina NovaSeq XPlus system (1.5B flow cell).

### Transcriptomics data analyses

The *Ostrinia nubilalis* reference genome (GCA_963855985.1) was indexed by STAR (2.7.11b) ^94^ using the RefSeq Genes annotation track (GCF_963855985.1) with rRNA genes removed (gene_biotype “rRNA”). Raw sequencing reads for all 64 samples were aligned to the genome by STARsolo in alignReads mode following BRB-seq recommendations, except *--outFilterScoreMinOverLread 0.3*, *--outFilterMatchNminOverLread 0.3*. Uniquely mapped reads that overlapped genes were counted using the STARsolo output. Sample Assignment of the total raw reads to individual samples was high, and 88.2% of the reads containing valid barcodes (n = 936,026,352 reads). The quality of the sequencing data was robust, with 96.9% of the bases in cell barcodes and UMIs and 94.8% of the bases in RNA reads having a score ≥Q30. In terms of alignment, 62.2% of the reads mapped uniquely to the reference genome, and 66.9% of these reads overlapped with unique gene features. Of the 17,376 total annotated non-rRNA genes in ECB, 15,155 genes were detected in at least one sample. To remove inconsistently expressed genes across sample timepoints and backgrounds, only genes with at least 0.50 counts per million (CPM) in 25% of samples were retained for further analyses. This resulted in 10,079 genes. All analyses were performed after converting gene counts to logarithmic space via the transformation Log_2_(CPM+1). Samples with <300,000 reads assigned to the genes were excluded from the analysis. Consequently, the number of raw reads assigned per sample ranged from 527,582 to 27,630,712 (mean: 9,582,975). Gene count tables were quantile normalized with the *voom* function from *limma* v3.32.5 ^95^. Significantly rhythmic (i.e., cycling) genes, differentially rhythmic, and differentially expressed genes were identified using a cosinor regression in LimoRhyde ^57^. Differences in amplitude, phase, and mesor (rhythm-adjusted mean) were further investigated for candidate circadian clock genes by *CircaCompare* ^96^. To control false discovery rate, *p* values were converted to *q-*values using the Benjamini and Hochberg (1995) method. Gene orthologs were identified by reciprocal protein BLAST ^97^ of *O. nubilalis* RefSeq proteins (GCF_963855985.1) and *D. plexippus* (GCF_009731565.1; see Table S1).

### CRISPR/Cas9 mutagenesis and mutant genotyping

CRISPR/Cas9 mutagenesis was performed as described previously ^62^. Briefly, early-stage embryos were injected with single sgRNA ribonucleoprotein (RNP) complexes targeting *per* (exon 4; see Table S2). DNA was extracted in a 1x DirectPCR tail lysis buffer (Viagen Biotech) containing 1 µg of Proteinase K (ThermoFisherScientific, USA) in a total volume of 100 µL. Samples were incubated for 16 hr at 56°C and 85°C for 25 min. Germline mutants harboring a frameshift mutation were verified by PCR and Sanger sequencing (Eton Bioscience).

Experimental individuals were genotyped by allele-specific PCR for known frameshift mutations (Family A-D; Table S1). To limit amplification of the wild-type allele, mutant allele-specific primers were designed with an additional mismatch in the first five bases of the 3’ end. In general, PCRs contained 1X GoTaq Master Mix (Promega), 0.1 µM of mutant allele-specific primers, 0.1 µM of locus-specific primers (positive control for DNA quality), and 1.5 µL of DNA extract in 20 µL.

To increase throughput, insects from Family E-F were genotyped by amplicon sequencing ^98^. Briefly, the first round of PCR with gene-specific primers was 20 cycles and gene-specific amplicons were diluted ten-fold before barcoding. Barcoded amplicons were purified using HighPrep PCR magnetic beads (MagBio Genomics, AC-60005) before Illumina preparation and sequencing (Amplicon-EZ service at Genewiz by Azenta). All oligos were synthesized by Integrated DNA Technologies and Eton Bioscience (Table S2).

Genomic reads were trimmed using TrimGalore to remove Illumina adapters and low-quality bases. CutAdapt v.3.7 ^99^ demultiplexed trimmed sequences by internal barcodes (-e 0.1). Due to overlapping paired-end reads, reads were merged by NGmerge v.0.3 ^100^ with modified settings (-v -d -m 50). Merged sequences smaller than 110 bases were removed by Seqtk v.75. HISAT2 ^101^ and aligned to a subset of the *O. nubilalis* reference genome (GCA_963855985.1) containing only the predicted amplicons. HISAT2 trimmed the 5’ linker and primer sequences from both 5’ and 3’ ends (--trim5 37 --trim3 37) of the merged reads, before aligning (-k 1 – score-min L,0,-0.6 --no-spliced-alignment). Alignments in BAM format were sorted and indexed by samtools v.1.2.0 ^102^. SNPs and small indels (<50 bp) were called using the GATK4 HaplotypeCaller ^103^ for females (--sample-ploidy 1) and males/unknown sex (--sample-ploidy 2). GVCFs were combined by *CombineGVCFs* and *GenotypeGVCFs* performed joint genotyping. *VariantFiltration* hard filtered variants by (QD < 2.0 || QUAL < 30 || MQ < 40.0). SNPs were selected and filtered to remove uninformative variants (AF <= 0.15 || AF >= 0.85).

VCFs were parsed into R by *vcfR* v.1.15.0 ^104^. For each sample genotype, allele balance was calculated as the proportion of allele counts for the alternate alleles. Heterozygous genotypes with an allele balance (AB) < 0.2 or AB > 0.8 were respectively filtered to homozygous for the reference or alternate allele ^105^. To conservatively remove false homozygotes, any genotypes with an allele depth (DP) < 5 were filtered to NA.

### Statistical Analyses

All statistical analyses were conducted in R^106^. The effects of genetic background and/or photoperiod on diapause incidence and cumulation pupation were quantified by a binomial logistic regression with a logit link. Linear regression evaluated the effects of period mutation on PDD-time and mass. To account for uneven sampling among families, square root transformed sum contrasts for family, period mutation, and LD photoperiod were used to understand how each factor level deviated from the overall mean. Differences in the distribution of adult eclosion were compared using Watson’s U^2^ test within the circular package^107^. All regression models were built in lme4 and used *Anova*() from the *car* package^108^ to determine the significance of main effects. Significant differences between group proportions and means were respectively evaluated by Wald’s Z test and Welch’s T test.

## Notes

https://github.com/daytonjn/period_ms_2024

### Competing Interest Statement

The authors have declared no competing interest.

