## Supplementary material for "The *period* gene alters daily and seasonal timing in *Ostrinia nubilalis*": Document S1

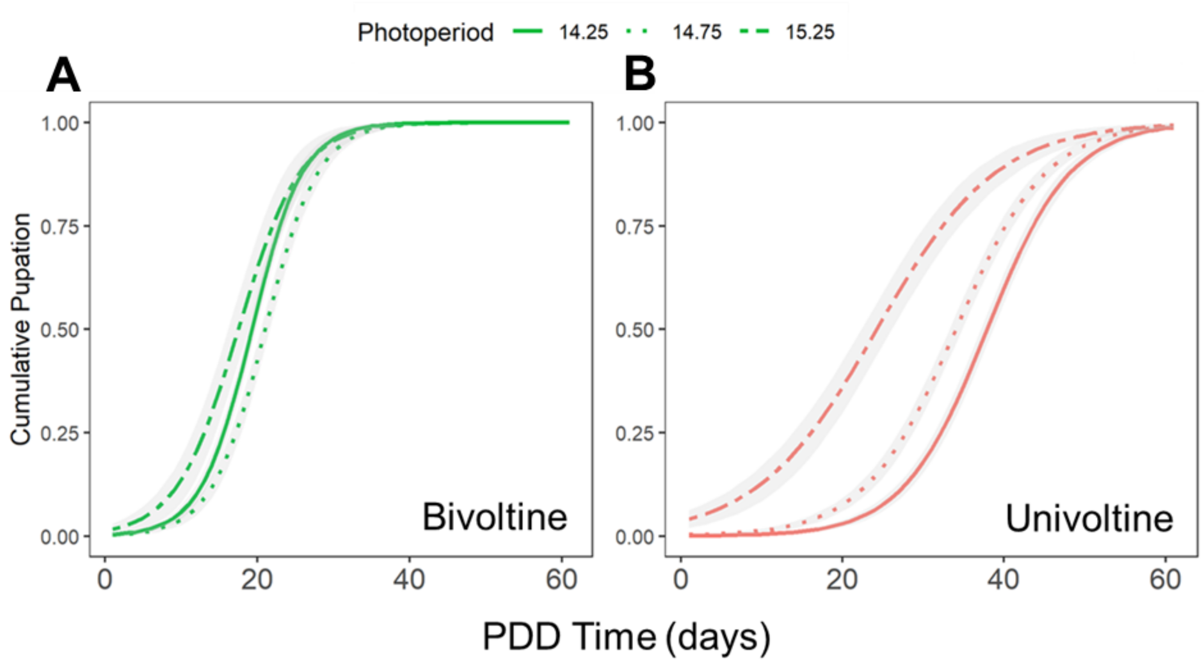


**Fig. S1. PDD time is altered by the diapause-inducing photoperiod.** Diapausing bivoltine (A) and univoltine (B) larvae were transferred to 16L:8D to track post diapause development (PDD) time. PDD time was significantly affected by photoperiod (binomial GLM; $\chi^2 = 123$, *df* = 1, $P < 2.2*10^{-16}$), ecotype ($\chi^2 = 1810$, *df* = 2, $P < 2.2*10^{-16}$) and their interaction ($\chi^2 = 125$, *df* = 2, $P < 2.2*10^{-16}$).

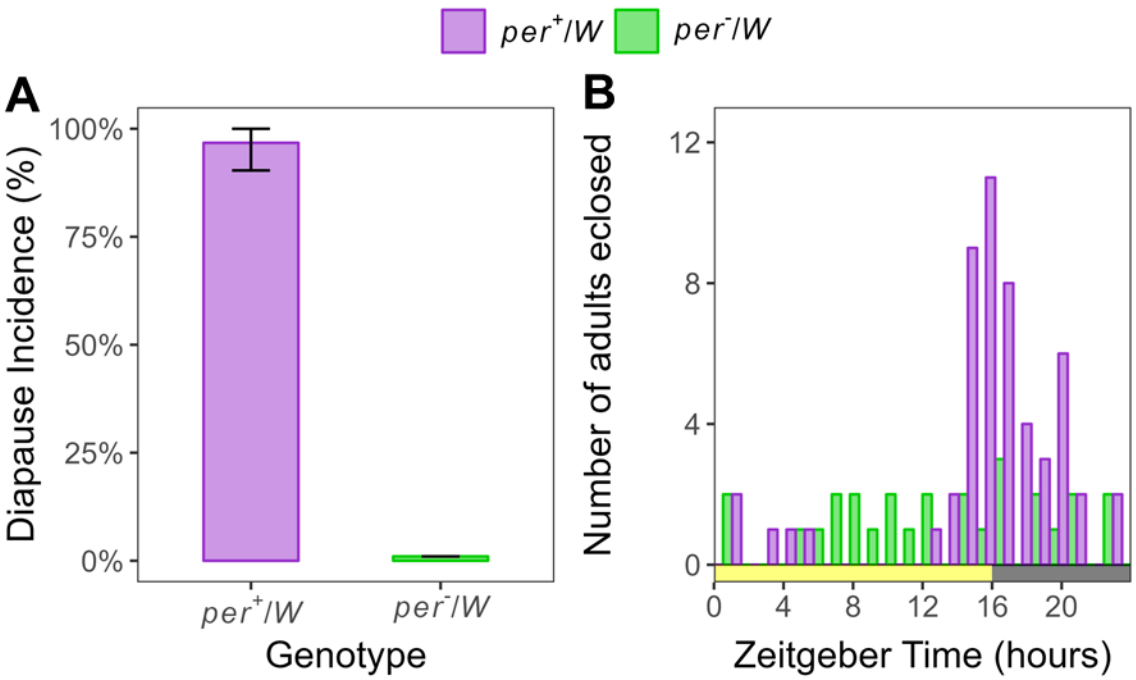


**Fig S2. *period* is essential for short-day recognition and daily timing.** (A) Diapause induction in larvae exposed to 12L:12D (23.5˚C). Compared to *per* wildtype ($per^{+/W}$) siblings (n = 74), diapause incidence was abolished in *per* hemizygous mutants ($per^{-/W}$) (n = 59; $\chi^2$ = 117.6, *df* = 1, $P < 0.001$). (B) Eclosion of female adults from their pupal case was pooled and binned into 1 hr intervals. Adults were entrained throughout their pupal stage to 16L:8D. Rhythmic eclosion of wildtype females (n = 53; Rayleigh's Z = 0.686, $P < 0.001$) was abolished in mutant *period* knockouts (n = 28; Z = 0.109, $P = 0.72$).

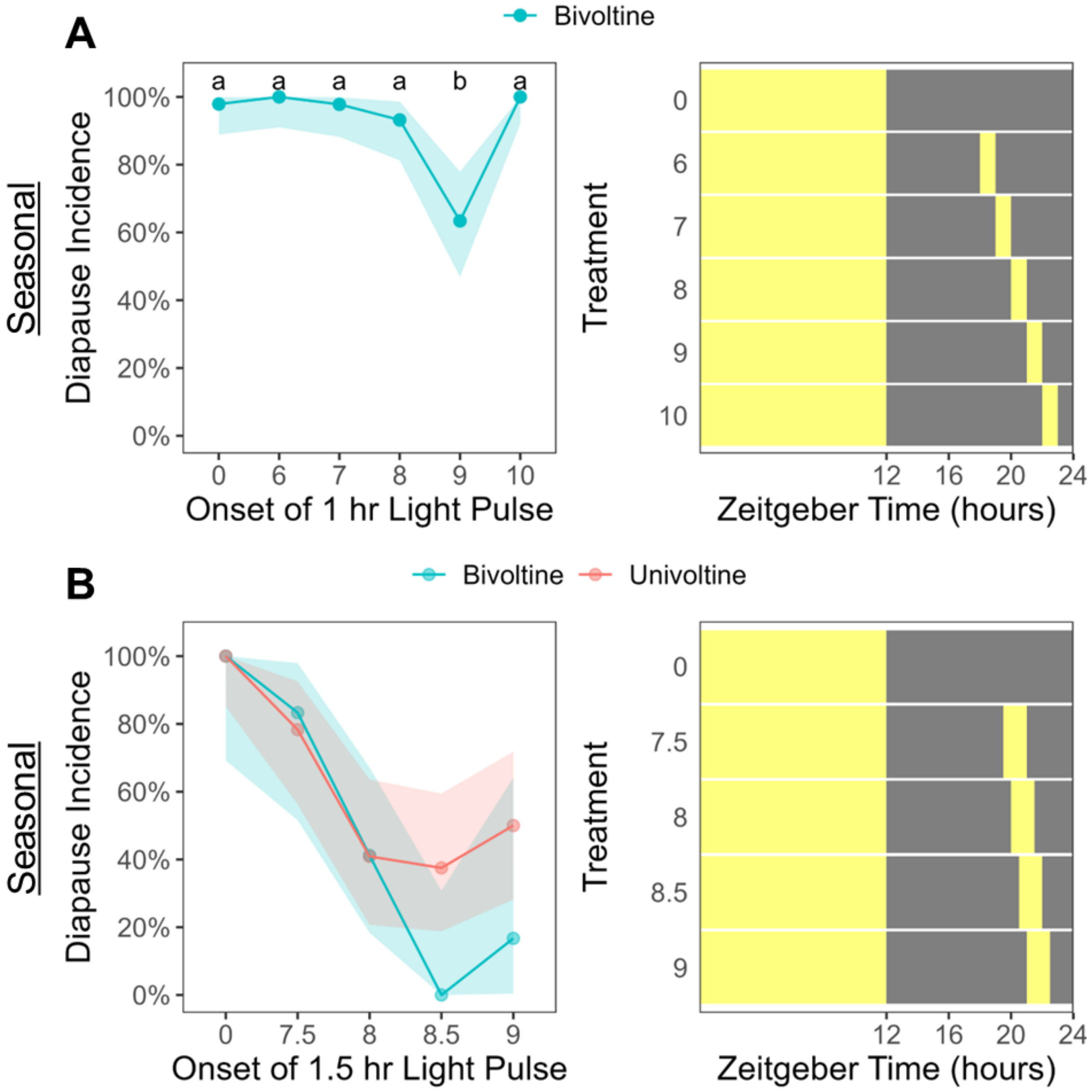


**Fig. S3. Night interruption disrupts diapause induction in *O. nubilalis*.** (A) Diapause induction of larvae reared in a 12L:12D photoperiod with a night-interrupting light pulse during the darkness (no pulse = 0). Letters denote significant differences between treatments (Z-test, $P < 0.05$). (A, Right) For each treatment, the onset and duration of the light pulse, in relation to the overall Zeitgeber time, is depicted with yellow (light) and black (dark) rectangles. (B) A binomial GLM estimated the effects of photoperiod treatment, ecotype, and their interaction on diapause incidence. Diapause incidence was significantly affected by photoperiod treatment ($\chi^2 = 65.3$, $df = 4$, $P < 2.2*10^{-13}$), but not by genetic background ($\chi^2 = 2.4$, $df = 1$, $P = 0.123$) or their interaction ($\chi^2 = 7.6$, $df = 4$, $P = 0.106$). Shading denotes 95% confidence intervals.

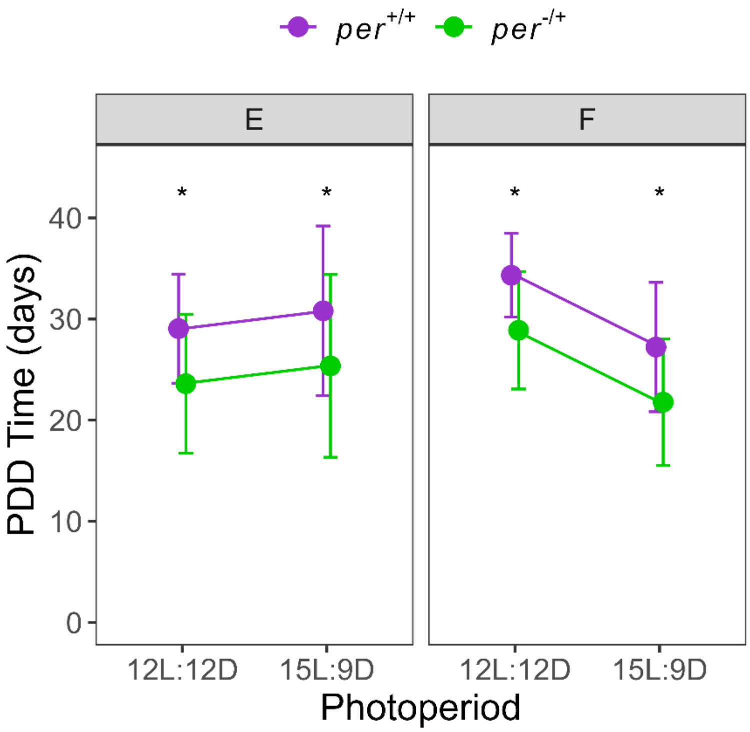


**Fig. S4. The effect of *per* gene dosage on PDD time is independent of the diapause-inducing photoperiod.** Diapausing larvae from 12L:12D and 15L:9D hr were transferred to 16L:8D to track post-diapause development (PDD) time. Although *per* gene dosage (two-way ANOVA: $F = 4.64$, $df = 1$, $P = 0.033$) and photoperiod ($F = 2.45$, $df = 1$, $P = 0.121$) influenced PDD time, there was no evidence for an interaction between *per* gene dosage and photoperiod ($F = 0.38$, $df = 1$, $P = 0.539$). Mean PDD time of families E and F are plotted with 95% confidence intervals. Asterisks denote significant differences in PDD time between genotypes, $P < 0.05$ (*).

**Table S1. GenBank accessions for *O. nubilalis* clock genes.** Orthologs were identified by reciprocal blast to *D. plexippus*.

| Gene | *O. nubilalis* | *D. plexippus* |
|---|---|---|
| *clock* | LOC135086749 | LOC116777439 |
| *bmal1* | LOC135075772 | LOC116770138 |
| *cycle* | LOC135086822 | LOC116778126 |
| *cry2* | LOC135082303 | LOC116765947 |
| *period* | LOC135086880 | LOC116777810 |
| *vrille* | LOC135084225 | LOC116775896 |
| *tim* | LOC135087922 | LOC116765334 |
| *pdp1-e* | LOC135086827 | LOC116778133 |
| *cwo* | LOC135088456 | LOC116765547 |
| *cry1* | LOC135074722 | LOC116774075 |

**Table S2. Oligos for sgRNA templates or PCR genotyping.** Bolded sequences denote the *O. nubilalis* sgRNA target sequence. Lowercase denotes linker forward and reverse linker primers to adhere barcodes from Liu et al. (2021).

| Oligo | Sequence |
|---|---|
| per_sgRNA1 | TTCTAATACGACTCACTATA**GGAGAAGAGGACCAAAGAGA**GTTTTAG AGCTAGA |
| JD43F | TCCAGGACTTCGGCTGAAAC |
| per_exon4R | CATCGCTGACCTCCTCCTTC |
| JD51R_x4_28del | TGCTTCTTCTTTTTGGGTCCTCT |
| JD55F_x4_90del | CGGAGAAGAGGACCAAAGAAG |
| per_mutx3_F | gtgaccaagttcatgctTTCATAGTGGCCCACGAGTAC |
| per_mutx3_R | gtcggagtcaacggattCATCGCTGACCTCCTCCTTC |
| per_mutx4_F | gtgaccaagttcatgctGTTATCCAGTTGCCTTCGCC |
| per_mutx4_R | gtcggagtcaacggattGGAGGTTACTGGTGGATGCA |
| pdfr_mutx3_F | gtgaccaagttcatgctGCCATCTCTGACTTCAGGACG |
| pdfr_mutx3_R | gtcggagtcaacggattACGTTGTTAAAGGAATTGCGCT |
| pdfr_mutx2_F | gtgaccaagttcatgctAAGCTCGATGACTTCAGGTGG |
| pdfr_mutx2_R | gtcggagtcaacggattACGGCTTTCGATTCCACAAAA |
| cipc_multi_F | gtgaccaagttcatgctCGGATCTGAACCGTCTGCG |
| cipc_multi_R | gtcggagtcaacggattTCGGAGGTTTGTTGTTGGAGT |
| tpi_multi_F | gtgaccaagttcatgctTGCATGCAACCGAAAGTTACA |
| tpi_multi_R | gtcggagtcaacggattAGTGACCGATCCTCCGTACT |
| tpi_multi_F | gtgaccaagttcatgctTGCATGCAACCGAAAGTTACA |
